# Distribution of Purines and Pyrimidines over miRNAs of Human, Gorilla and Chimpanzee

**DOI:** 10.1101/208405

**Authors:** Jayanta Kumar Das, Pabitra Pal Choudhury, Adwitiya Chaudhuri, Sk. Sarif Hassan, Pallab Basu

**Author notes:** Equal contributor.

## Abstract

Meaningful words in English need vowels to break up the sounds that consonants make. The Nature has encoded her messages in RNA molecules using only four alphabets A, U, C and G in which the nine member double-ring bases (adenine (A) and Guanine (G)) are purines, while the six member single-ring bases (cytosine (C) and uracil (U)) are pyrimidines. Four bases A, U, C and G of RNA sequences are divided into three kinds of classifications according to their chemical properties. One of the three classifications, the *purine-pyrimidine* class is important. In understanding the distribution (organization) of purines and pyrimidines over some of the non-coding regions of RNA, all miRNAs from three species of Family Hominidae (namely human, gorilla and chimpanzee) are considered. The distribution of purines and pyrimidines over miRNA shows deviation from randomness. Based on the quantitative metrics (fractal dimension, Hurst exponent, Hamming distance, distance pattern of purine-pyrimidine, purine-pyrimidine frequency distribution and Shannon entropy) five different clusters have been made. It is identified that there exists only one miRNA in human *hsa-miR-6124* which is purely made of purine bases only.

**AMS Subject Classification:** 92B05 & 92B15

## Introduction

The monomer units for forming the nucleic acid polymers deoxyribonucleic acid (DNA) and ribonucleic acid (RNA) are nucleotides which are grouped into three different classes based on their chemical properties, i.e., purine group R={A,G} and pyrimidine group Y={C,T/U}; amino group M={A,C} and keto group K={G,T/U}; strong H-bond group S={C,G} and weak H-bond group W={A,T/U} [1]. There are two kinds of nitrogen-containing bases-Purines and Pyrimidines, first isolated from hydrolysates of nucleic acids, were identified using classical methods of organic chemistry. An important contribution was made by Emil Fischer who must be credited with the earliest synthesis of purines (1897) [2]. Purines consist of a nine member double-ring (containing carbon and nitrogen) fused together, where as pyrimidines have a six member single-ring comprising of carbon and nitrogen. [3, 4]. The organization of purine-pyrimidine bases over RNA is crucial. Here we intend to understand the organization of the two chemical bases purine and pyrimidine over over some of the non-coding RNAs, *microRNA*.

MicroRNAs (abbreviated miRNAs) contain about 18-25 ribonucleotides that can play important gene regulatory roles by pairing to the messages of protein-coding genes, to specify messenger RNA (mRNA) cleavage or repression of productive translation [5, 6, 7]. miRNA genes are one of the more abundant classes of regulatory genes in animals, estimated to comprise between 0.5 and 1 percent of the predicted genes in worms, flies, and humans, raising the prospect that they could have many more regulatory functions than those uncovered to date [8,9]. The main function of miRNAs is to down-regulate gene expression [10]. A given miRNA may have hundreds of different mRNA targets, and a given target might be regulated by multiple miRNAs [11, 12, 13, 14, 15]. It is important to identify the miRNA targets accurately. miRNAs control gene expression by targeting mRNAs and triggering either translation repression or RNA degradation [16,17]. Their aberrant expression may be involved in various human diseases, including cancer [18, 19, 20, 21, 22, 23, 24]. miRNA regulatory mechanisms are complex and there is still no high-throughput and low-cost miRNA target screening technique [25, 26, 27]. It is an well known fact that each miRNA is potentially able to regulate around 100 or more targeted mRNAs and 30% of all human genes are regulated by miRNAs [28].

In this article an attempt has been made to decipher the pattern of organization of purines and pyrimidines distributions over the miRNAs of the three species human, gorilla and chimpanzee. We desire to understand how the purine/pyrimidine bases are organized over the sequence and how much distantly the purine/pyrimidine bases can be placed over the sequence. Which one of these two type of chemical bases purine or pyrimidine dominates other in terms of their frequency density over the sequence is one of our prime aims to comprehend. A simple binomial distribution (i.e. location independent occurance of the bases) fails to describe the observed variation of Purine and Pyrimidine. This encourages us to look for further patterns. We investigate, the self-organization of the purine and pyrimidine bases for all the miRNAs of the three species human, gorilla and chimpanzee through the fractal dimension of the indicator matrix. The auto correlation of purine-pyrimidine bases over the miRNAs through the parameter Hurst exponent are determined and found many of the miRNAs have identical auto correlation even if their purine-pyrimidine organization is different. All the miRNAs are compared about their nearness based on their purine-pyrimidine distribution, Hamming distance is employed among all the miRNAs in understanding the nearness of purine-pyrimidine organization. The purine-pyrimidine distance patterns including the frequency distribution have been found for all the miRNAs for all three species. All possible distinct pattern of frequency distribution are determined for all the miRNAs of both the species. Here we wish to bring attention to the reader that through our investigation, the miRNA h1291, made of only purine bases is identified. There is no miRNA (human, gorilla and chimpanzee) which is absolutely made of pyrimidines.

## Materials and Methods

### Datasets

From the MiRBase (a miRNA database: http://www.mirbase.org/) [29], from the family Hominidae, total of 2588 mature miRNAs of human, 357 mature miRNAs of gorilla and 587 mature miRNAs of chimpanzee are taken. Each miRNA of human, gorilla and chimpanzee are encoded as numbers starting from *h*1 to the total number of sequences h2588 for miRNAs of human and same has been made for miRNAs of gorilla *g*1 to *g*357 and for miRNAs of chimpanzee *p*1 to *p*587 (*Additional file 1*). We then transform the miRNAs sequences (A, U, C, G) into binary sequences (1’s and 0’s) according to the following rules:

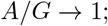

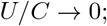

That means purine and pyrimidine nucleotide bases are encoded as 1 and 0 respectively into the transformed binary sequences of miRNAs. Therefore, presently we have three datasets of binary sequences of the three species human, gorilla and chimpanzee. All the computational codes are written in *Matlab R2016a* software.

### Fractal Dimension of Indicator Matrices

Here we shall encode each binary sequences into its indicator matrices [30, 31]. It is noted that there are several other technique for finding fractal dimension anf self organization structure of DNA sequences [32,33]. Consider a set S = {0,1} and an indicator function *f*: {0,1} × {0,1} ⟷ {0,1} is defined as for all (*x, y*) ∈ **S** × **S**,

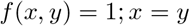

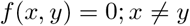

This indicator function can be used to obtain the binary image of the binary sequence as a two dimensional dot-plot. The binary image obtained by this indicator matrix can be used to visualize the distribution of ones and zeros within the same binary sequence and some kind of auto-correlation between the ones and zeros of the same sequence. It can be easily drawn by assigning a black dot to 1 and a white dot to 0. An example of indicator matrix is shown in Fig. 1 for the binary sequence *Hsa* – *miR* – 576 – 3*pMIM AT*0004796: 1111010111111100111100. From the indicator matrix, we can have an idea of the “fractal-like” distribution of ones and zeros (purines and pyrimidines). The fractal dimension for the graphical representation of the indicator matrix plots can be computed as the average of the number *p*(*n*) of 1 in the randomly taken *n* × *n* minors of the *N* × *N* indicator matrix. Using *p*(*n*), the fractal dimension (FD) is defined as

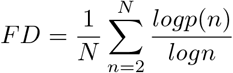

**Figure 1.**
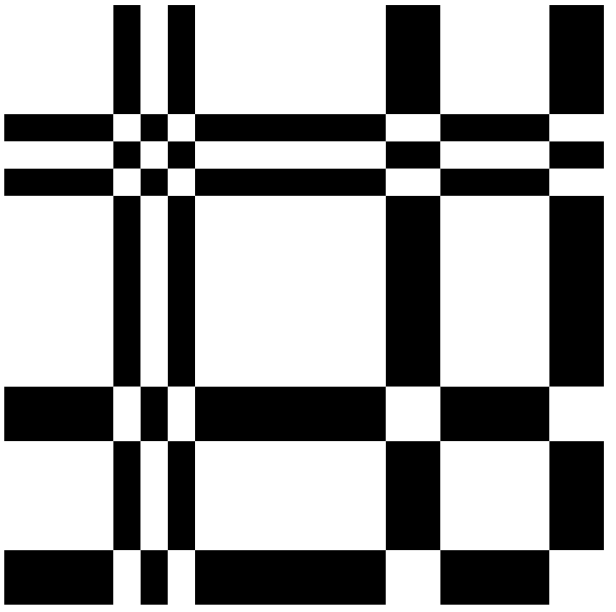
Indicator matrix for the binary sequence *Hsa* – *miR* – 576 – 3*pMIM AT*0004796: 1111010111111100111100

### Hurst Exponent of Binary Sequences

The Hurst Exponent (HE) deciphers the long term memory of time series, i.e. the autocorrelation of the time series [34, 35, 36]. A value of 0 < *HE* < 0.5 indicates a time series with negative autocorrelation and a value of 0.5 < *HE* < 1 indicates a time series with positive autocorrelation. A value of *HE* = 0.5 indicates a true random walk, where it is equally likely that a decrease or an increase will follow from any particular value with higher values indicating a smoother trend and less roughness.

The Hurst exponent *HE* of a binary sequence {*x*_*n*_} is defined as 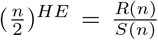 where 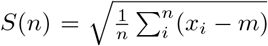 and *R*(*n*) = *maxY*(*i,n*) − *minY*(*i,n*); 1 ≤ *i* ≤ *n* where 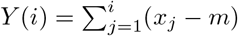 and 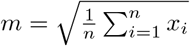

### Hamming Distance of Binary Sequences

The Hamming Distance (*HD*) between two binary strings is the number of bits in which they differ [37, 38, 39]. Since length of the miRNAs might differ and hence a special care has been taken into consideration. Suppose there are two miRNAs *S_m_* and *S_n_* of length *m* and *n* respectively (*n* > *m*), then

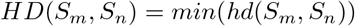

where *S*_*n*_ of length *n* window is sliding over *S*_*m*_ from the left alignment to the right alignment and each time hamming distance (hd) is calculated, and finally minimum *hd* value is taken as hamming distance *HD* of two binary sequences.

For example, take two binary sequences *S*_*m*_ = 010100 and *S_n_* = 1101, now sliding of *S*_*n*_ over *S*_*m*_ of length 4, from left to right alignment of these two sequences, we find the hamming distances are hd(**0101**00,**1101**)=1, hd(0**1010**0,**1101**)=3, hd(01**0100**,**1101**)=2, therefore we take *HD* = 1 (minimum) of these two binary sequences. Finding the minimum hamming distance of the two binary sequences says about the maximum similarity of two sequences over the distribution of purines and pyrimidines. The minimum value of *HD* = 0 when the pattern of length *min*(*m, n*) of two binary sequences of miRNAs are exactly identical i.e. similar distribution of purines and pyrimidines over the miRNAs of the two sequences and the maximum value of *HD* = *min*(*m,n*) when the pattern of length *min*(*m,n*) of two binary sequences of miRNAs are exactly opposite i.e. completely dissimilar distribution of purines and pyrimidines over miRNAs two sequences.

### Distance pattern of purine and pyrimidine over miRNAs

Here we are exploring the distance pattern of purines bases across the miRNAs human, gorilla and chimpanzee. How sparsely (closely) purine bases are placed over the miRNAs. So we found the distance (gap) between purine bases to the immediate next purine base over the miRNA sequences.

For example, take a transformed binary sequence *S*_*m*_ = 110100111000001, where 1 indicates the purines bases and 0 indicates pyrimidine bases in the sequence. From left to right the positions of 1’s and 0’s in serial is shown below. Now, form the distribution of 1’s, we find the purine distances at 1 (two consecutive 1’s at a distance of 1: 11), 2 (two consecutive 1’s at a distance of 2: 101), 3 (two consecutive 1’s at a distance of 3: 1001) and 6 (two consecutive 1’s at a distance of 6: 1000001). So, the distance pattern of purines over the sequence is [1 2 3 6] in order.

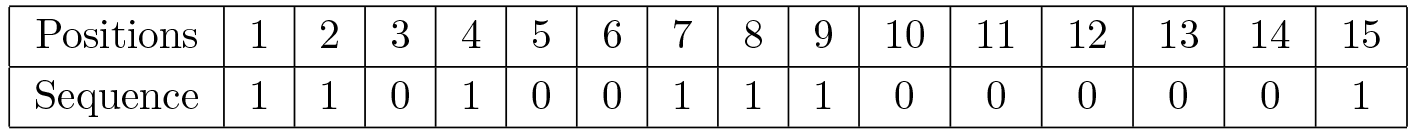

|  |  |  |  |  |  |  |  |  |  |  |  |  |  |  |  |
| --- | --- | --- | --- | --- | --- | --- | --- | --- | --- | --- | --- | --- | --- | --- | --- |
| Positions | 1 | 2 | 3 | 4 | 5 | 6 | 7 | 8 | 9 | 10 | 11 | 12 | 13 | 14 | 15 |
| Sequence | 1 | 1 | 0 | 1 | 0 | 0 | 1 | 1 | 1 | 0 | 0 | 0 | 0 | 0 | 1 |

Similar to the distance pattern of purine, the distance pattern of pyrimidine bases (0’s) across the miRNAs also can be determined. The distance pattern of pyrimidines of the above sequence is [1 2 4] in order. Further the distance pattern of pyrimidines [1 **2** 4] of a miRNA opens up a fact that there is at least one 1(= **2** − 1) and at least one 3(= **4** − 1) length purine blocks present in the miRNA. In the similar way, a distance pattern of purine triggers the presence of pyrimidine blocks in miRNAs.

### Shannon entropy of miRNAs

The Shannon entropy (SE) is defined as the entropy of a Bernoulli process with probability *p* of one of two values [40, 41, 42]. It is defined as

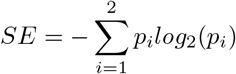

Here 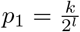 and 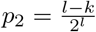; here *l* is length of the binary string and *k* is the number of 1s in the binary string of length *l*.

The *Shannon entropy* is a measure of the uncertainty in a binary string. To put it intuitively, suppose *p* = 0. At this probability, the event is certain never to occur, and so there is no uncertainty at all, leading to an entropy of 0. If *p* = 1, the result is again certain, so the entropy is 0 here as well. When *p* = 1/2, the uncertainty is at a maximum; if one were to place a fair bet on the outcome in this case, there is no advantage to be gained with prior knowledge of the probabilities. In this case, the entropy is maximum at a value of 1 bit. Intermediate values fall between these cases; for instance, if *p* = 1/4, there is still a measure of uncertainty on the outcome, but one can still predict the outcome correctly more often than not, so the uncertainty measure, or entropy, is less than 1 full bit.

## Results and Classifications of miRNAs

### Deviation from randomness

A simple random binomial (p,q) model [43], where each entries can either be purine (or pyrimidine) with probability *p* (or *q* = 1 − *p*) fails to address the distribution of purine or pyrimidine over miRNAs. If we calculate the mean of the distribution from the sample and calculate the probability *p*, and if we calculate the probability *p* from the variance then we get different values. In all three species, the expected variances are significantly smaller than what we would have expected from the binomial distribution.

In human *p* = 0.509, chimpanzee *p* = 0.505, gorilla *p* = 0.514 from mean. Expected std = 2.327 (human),2.323 (chimpanzee), 2.322 (gorilla). Sample std = 3.35 (human), 2.78 (chimpanzee), 2.58 (gorilla).

### Classification Based on FDs of Indicator Matrices

The fractal dimension for each binary sequence of human, gorilla and chimpanzee miRNAs as entire data is given in the *Additional file 2 (sheet-1)*. Based on the fractal dimension, we have made classifications for the all the datasets of human, gorilla and chimpanzee. There are 10 clusters of miRNAs for the species human, gorilla and chimpanzee as shown in Table 1. The fractal dimensions including the histograms of all the miRNAs of human, gorilla and chimpanzee are plotted in the Fig. 2. Also a normal distribution fitting is also made as shown in Fig.3. The detail members (miRNAs) of the clusters for human, gorilla and chimpanzee are given in the *Additional file 2 (sheet-2)*.

**Table 1.** Clusters based on Fractal dimension of miRNAs of Human, Gorilla and Chimpanzee.

| Cluster | Human |  | Gorilla |  | Chimpanzee |  |
| --- | --- | --- | --- | --- | --- | --- |
|  | Frequency | Center | Frequency | Center | Frequency | Center |
| 1 | 48 | 1.529 | 124 | 1.5683 | 19 | 1.5451 |
| 2 | 731 | 1.568 | 151 | 1.5995 | 255 | 1.5771 |
| 3 | 931 | 1.607 | 45 | 1.6308 | 189 | 1.6092 |
| 4 | 394 | 1.646 | 13 | 1.662 | 54 | 1.6413 |
| 5 | 222 | 1.685 | 12 | 1.6932 | 37 | 1.6734 |
| 6 | 137 | 1.724 | 7 | 1.7245 | 12 | 1.7055 |
| 7 | 80 | 1.763 | 3 | 1.7557 | 12 | 1.7375 |
| 8 | 26 | 1.802 | 0 | 1.7869 | 4 | 1.7696 |
| 9 | 14 | 1.841 | 0 | 1.8182 | 2 | 1.8017 |
| 10 | 5 | 1.88 | 2 | 1.8494 | 3 | 1.8338 |

**Figure 2.**
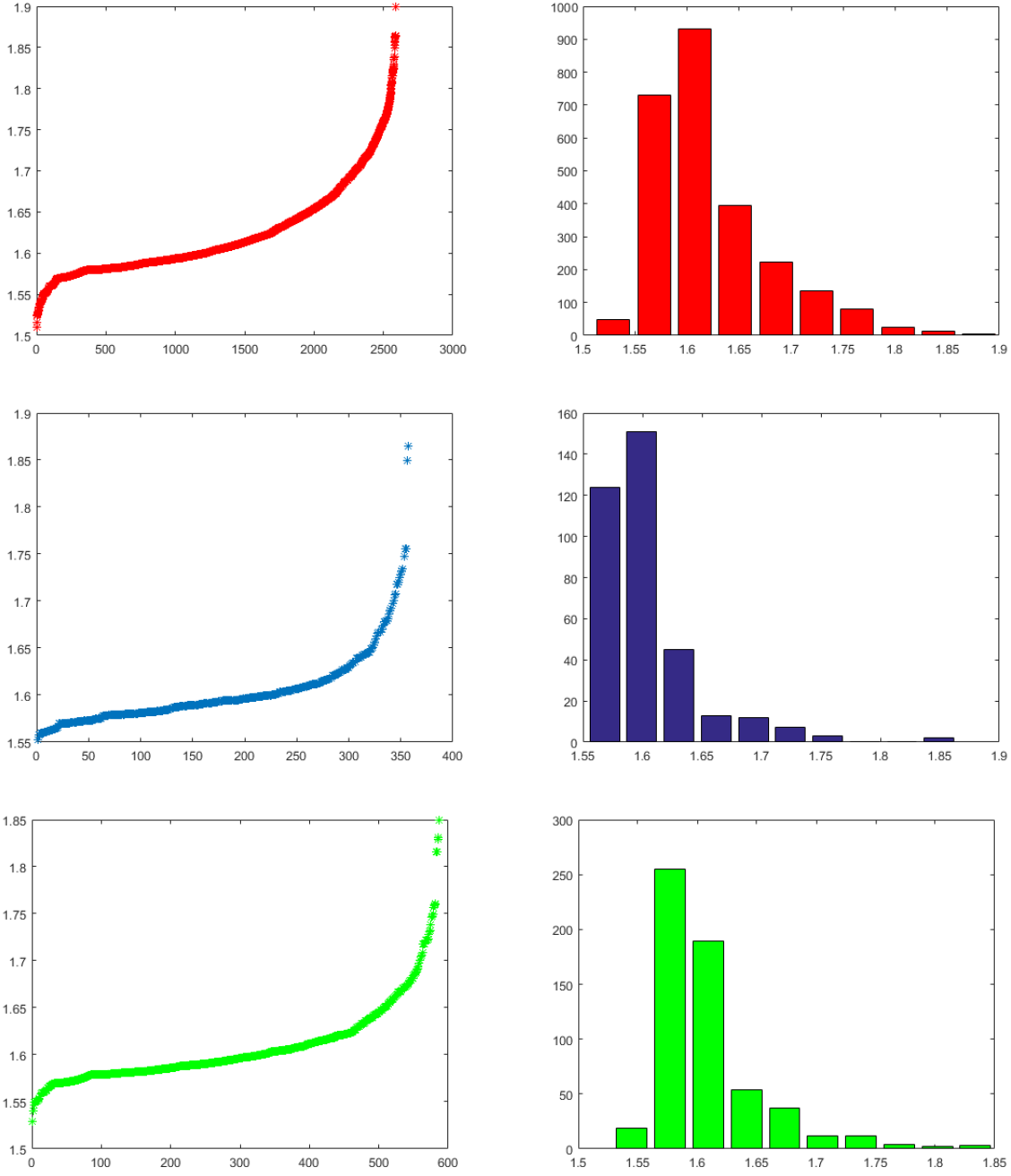
Histograms of fractal dimensions of miRNAs of Human, Gorilla and Chimpanzee from top to bottom respectively.

**Figure 3.**
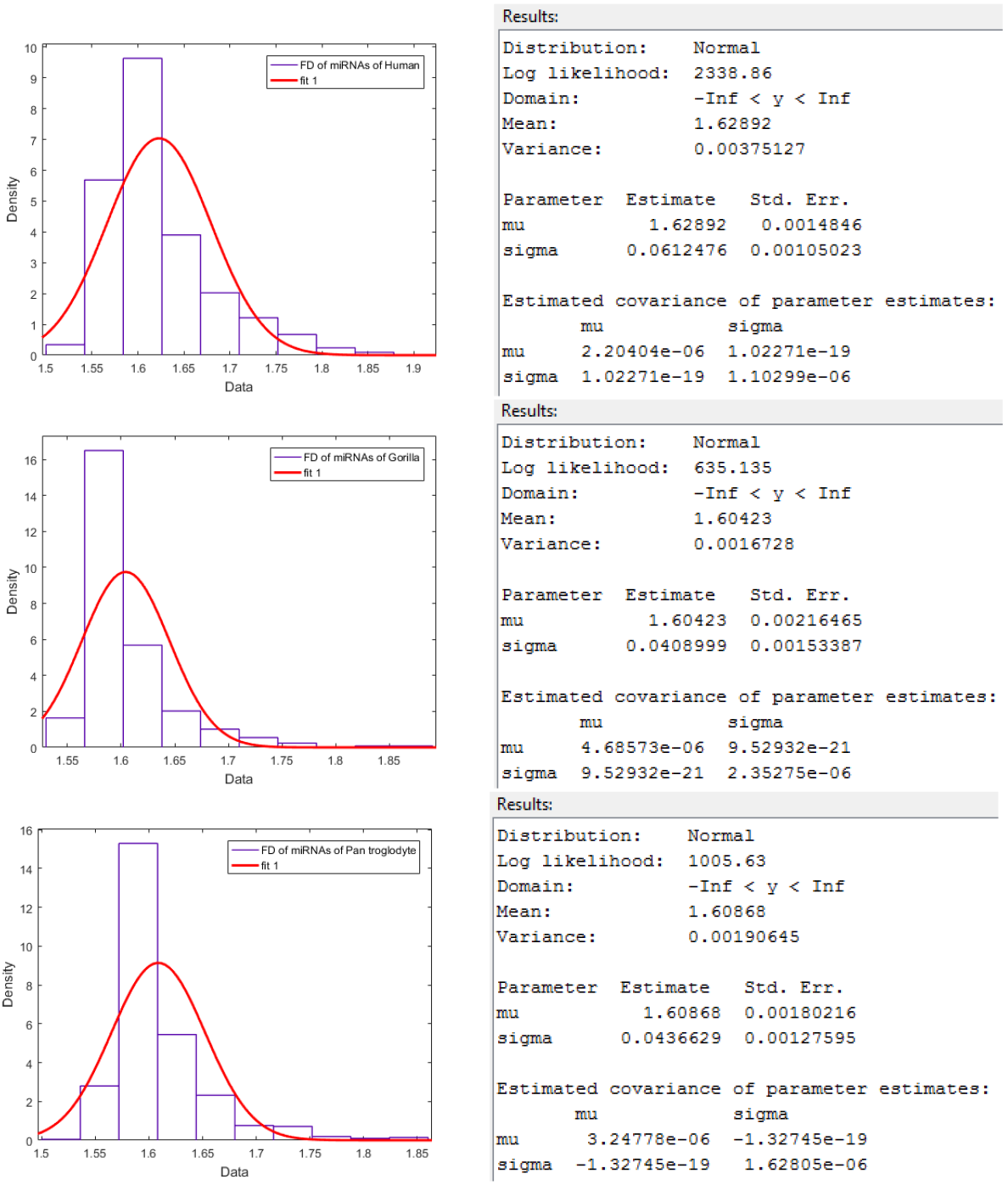
Normal distribution fitting over FDs of miRNAs of Human, Gorilla and Chimpanzee from top to bottom respectively.

It is observed that the FD of miRNAs of human lies in the interval [1.51,1.9] and the largest cluster(center at 1.60) contains 931 miRNAs whereas the FD of miRNAs of gorilla lies in the interval [1.55,1.864] which is contained in the interval [1.51,1.9] and the FD of miRNAs of chimpanzee lies in the interval [1.54,1.83] and the largest cluster (center at 1.58) contains 255 miRNAs. The largest cluster (center at 1.60) of miRNAs of gorilla contains 151 members. The centers of all the three largest clusters of miRNAs of human, gorilla and chimpanzee are approximately same which is a reflection of the fact that all these three species of the family *Hominidae* belong to the same family and are evolutionarily close. It is noted that there is no miRNAs of gorilla whose FD lies in between 1.76 and 1.84 whereas there are approximately 72 miRNAs of human and 7 miRNAs of chimpanzee whose FD lies in the said interval. There are clusters (for human, gorilla and chimpanzee) with largest centers among the other centers of the clusters contain 5, 2 and 3 members respectively.

### Classification Based on HEs

For every sequence of miRNA of human, gorilla and chimpanzee, Hurst exponent are determined ( *Additional file 3 (sheet-1)*) and then a classification is made based on the computed Hurst exponent for miRNAs of human, gorilla and chimpanzee, which is shown in Table 2. There are 10 clusters for all the species human, gorilla and chimpanzee miRNAs as shown in Table 2. The Hurst exponents and the histograms of all the miRNAs of human and gorilla are plotted in the Fig. 4. Also a normal distribution fitting is also made as shown in Fig.5. The detail members (miRNAs) of the clusters for human, gorilla and chimpanzee are given in the *Additional file 3 (sheet-2)*. The HE of miRNAs of human lies in the interval [0.2659, 0.9639] and the largest cluster(center at 0.7197) contains 671 miRNAs whereas the HE of miRNAs of gorilla lies in the interval [0.375, 0.964] and HE of miRNAs of chimpanzee in the interval [0.2745,0.9589]. The largest cluster (center at 0.64) of miRNAs of gorilla contains 69 members and the same(center at 0.7194) contains 134 miRNAs of chimpanzee. The centers of the HE clusters in the case human and chimpanzee are close enough where as the center of the largest cluster of miRNAs of gorilla is significantly different from other two species unlike FD as stated in the above section. It interprets basically the long range autocorrelations of miRNAs of gorilla is different significantly from the miRNAs of human and chimpanzee. It also shows the evolutionary closeness of human and chimpanzee in comparison with other two possible pairs (chimpanzee, gorilla) and (human, gorilla).

**Table 2.** Clusters based on Hurst exponent of miRNAs of Human, Gorilla and Chimpanzee.

| Cluster | Human |  | Gorilla |  | Chimpanzee |  |
| --- | --- | --- | --- | --- | --- | --- |
|  | Frequency | Center | Frequency | Center | Frequency | Center |
| 1 | 3 | 0.3009 | 7 | 0.4043 | 1 | 0.3087 |
| 2 | 11 | 0.3707 | 13 | 0.4632 | 4 | 0.3772 |
| 3 | 50 | 0.4405 | 28 | 0.5221 | 17 | 0.4456 |
| 4 | 173 | 0.5103 | 50 | 0.581 | 40 | 0.5141 |
| 5 | 430 | 0.5801 | 69 | 0.64 | 103 | 0.5825 |
| 6 | 600 | 0.6499 | 62 | 0.6989 | 123 | 0.651 |
| 7 | 671 | 0.7197 | 61 | 0.7578 | 134 | 0.7194 |
| 8 | 405 | 0.7895 | 30 | 0.8167 | 87 | 0.7879 |
| 9 | 168 | 0.8593 | 12 | 0.8756 | 57 | 0.8563 |
| 10 | 76 | 0.9291 | 25 | 0.9345 | 21 | 0.9248 |

**Figure 4.**
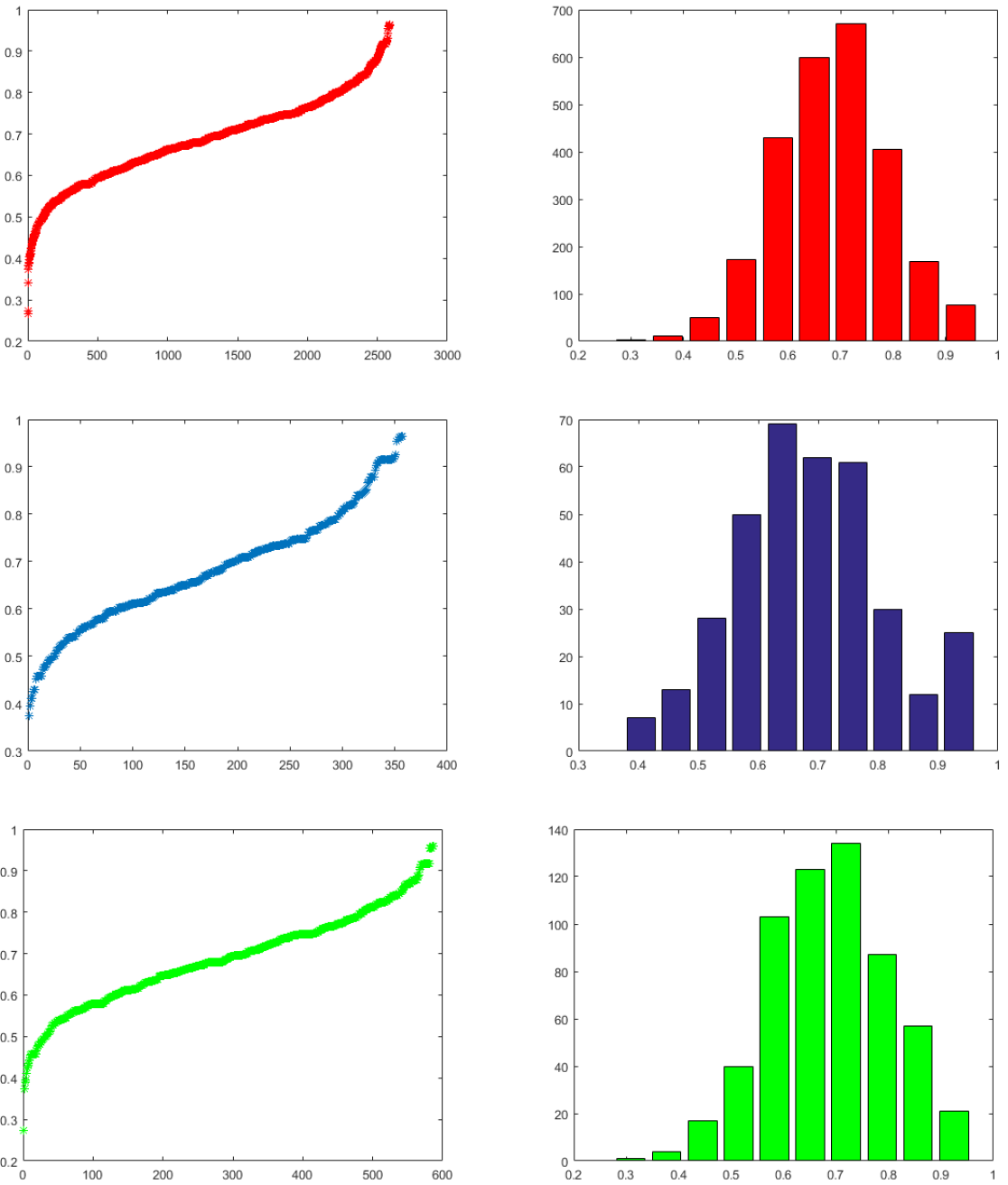
Histograms of Hurst exponents of miRNAs of Human, Gorilla and Chimpanzee from top to bottom respectively.

**Figure 5.**
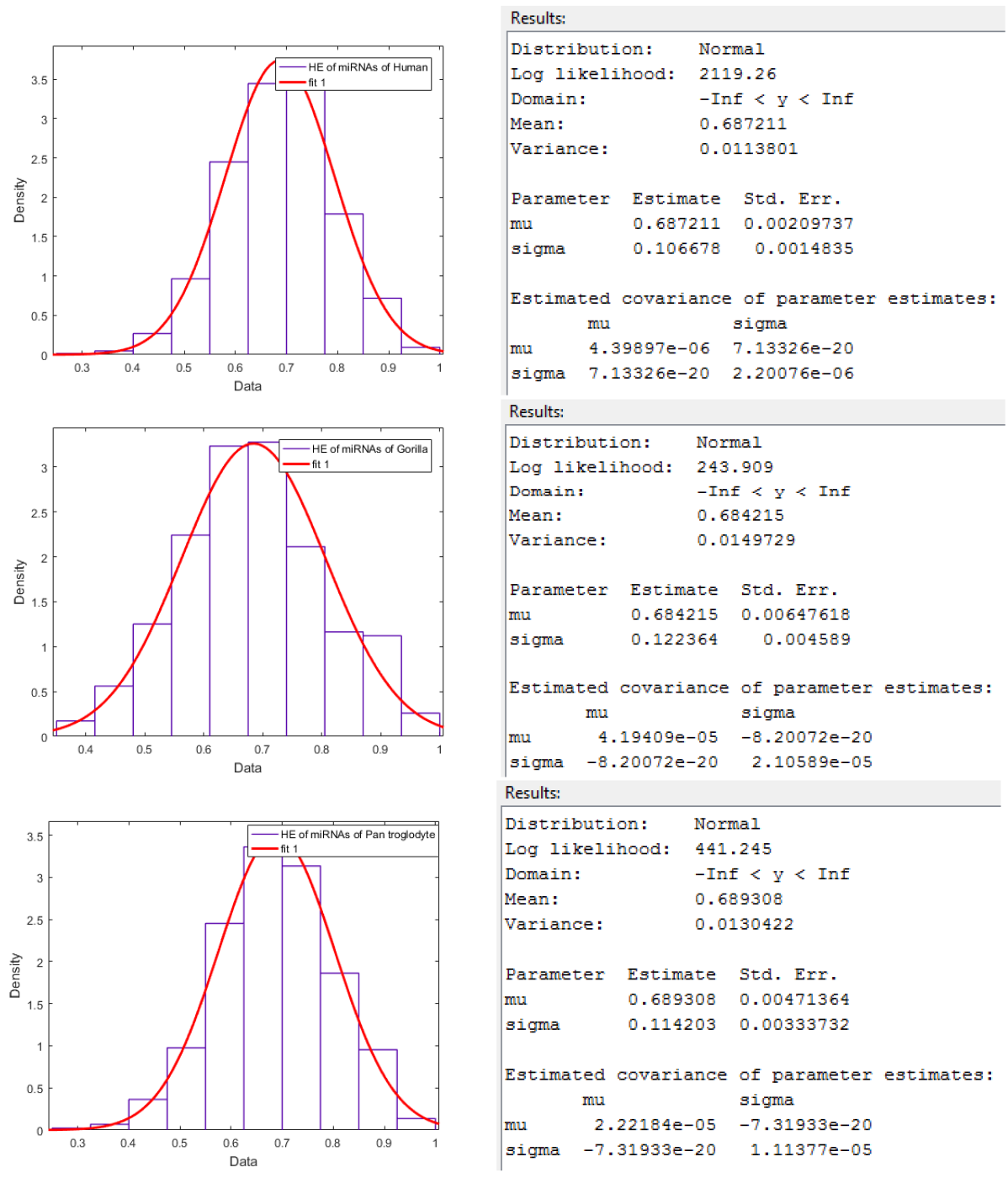
Normal distribution fitting over HEs of miRNAs of Human, Gorilla and Chimpanzee from top to bottom respectively.

### Classification Based on HDs

The detail pairs of miRNAs based on Hamming distances of human, gorilla and chimpanzee are given in the *additional file-4 (sheet-1)*, *additional file-4 (sheet-2)* and *additional file-4 (sheet-3)* respectively. We then form classes of pairs of the binary strings (miRNAs) of the three species human, gorilla and chimpanzee based on Hamming distances 0 to 21 as shown in Table 3. The bar plots with pie plots of these class frequencies are also given the Fig. 6.

**Table 3.** Clusters based on Hamming distance of miRNAs of Human, Gorilla and Chimpanzee. Here number of pairs for each hamming distance is shown, the details cal be seen from Additional files-4.

| Distance | Human | Gorilla | Chimpanzee |
| --- | --- | --- | --- |
|  | Number of pairs | Number of pairs | Number of pairs |
| 0 | 3768 | 557 | 763 |
| 1 | 2602 | 206 | 330 |
| 2 | 10620 | 244 | 572 |
| 3 | 39956 | 436 | 1264 |
| 4 | 119160 | 1154 | 4052 |
| 5 | 276592 | 3274 | 10996 |
| 6 | 505864 | 7370 | 23074 |
| 7 | 760868 | 14014 | 38752 |
| 8 | 960910 | 19714 | 51488 |
| 9 | 1042752 | 23076 | 56794 |
| 10 | 971648 | 20408 | 53210 |
| 11 | 781550 | 15476 | 41256 |
| 12 | 549660 | 9930 | 28722 |
| 13 | 339810 | 5844 | 17082 |
| 14 | 186330 | 3144 | 9332 |
| 15 | 89458 | 1444 | 4244 |
| 16 | 37768 | 720 | 1728 |
| 17 | 13404 | 284 | 646 |
| 18 | 3918 | 144 | 200 |
| 19 | 900 | 6 | 54 |
| 20 | 174 | 2 | 8 |
| 21 | 32 | 2 | 2 |

**Figure 6.**
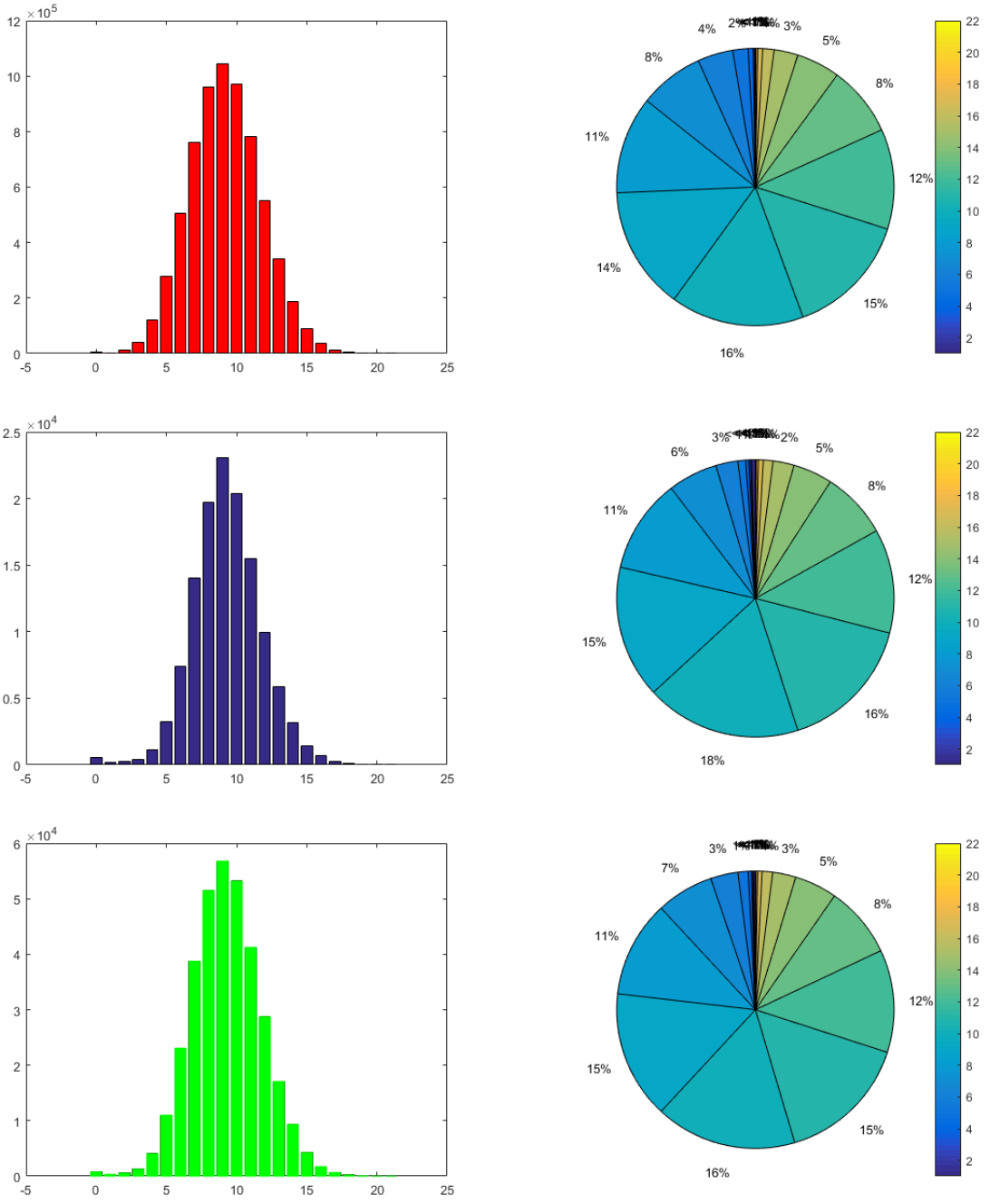
Bar plots and Pie plots of miRNAs of Human, Gorilla and Chimpanzee based on hamming distances from top to bottom respectively.

For all the three cases of miRNAs of human, gorilla and chimpanzee, none of the clusters with Hamming distances from 0 to 21 is empty. It is also seen that the largest clusters with HD 9 for miRNAs of human, gorilla and chimpanzee contain 1042752, 23076 and 56794 pairs respectively. It interprets that the arrangement of the purine and pyrimidine bases for most of miRNAs of human, gorilla and chimpanzee are differed by 9 bases only.

### Classification Based on Distance Pattern of Purine and Pyrimidine

For all the miRNAs of human, gorilla and chimpanzee, the distance patterns between purine bases to the next immediate purine bases are obtained. There are 187, 45 and 64 clusters based on unique distinct patterns of purine bases distance (gap) of miRNAs of human, gorilla and chimpanzee respectively *Additional file-5 (sheet-1,2,3)*. For an example, in the cluster 13 of miRNA of human, the pattern of purine distances in the miRNAs is [1 2 3 4 12] which is interpreted as there are purine bases which are 1, 2, 3, 4 and 12 bases apart. The bar plot of number of miRNAs for the three species human, gorilla and chimpanzee are plotted in Fig. 7.

**Figure 7.**
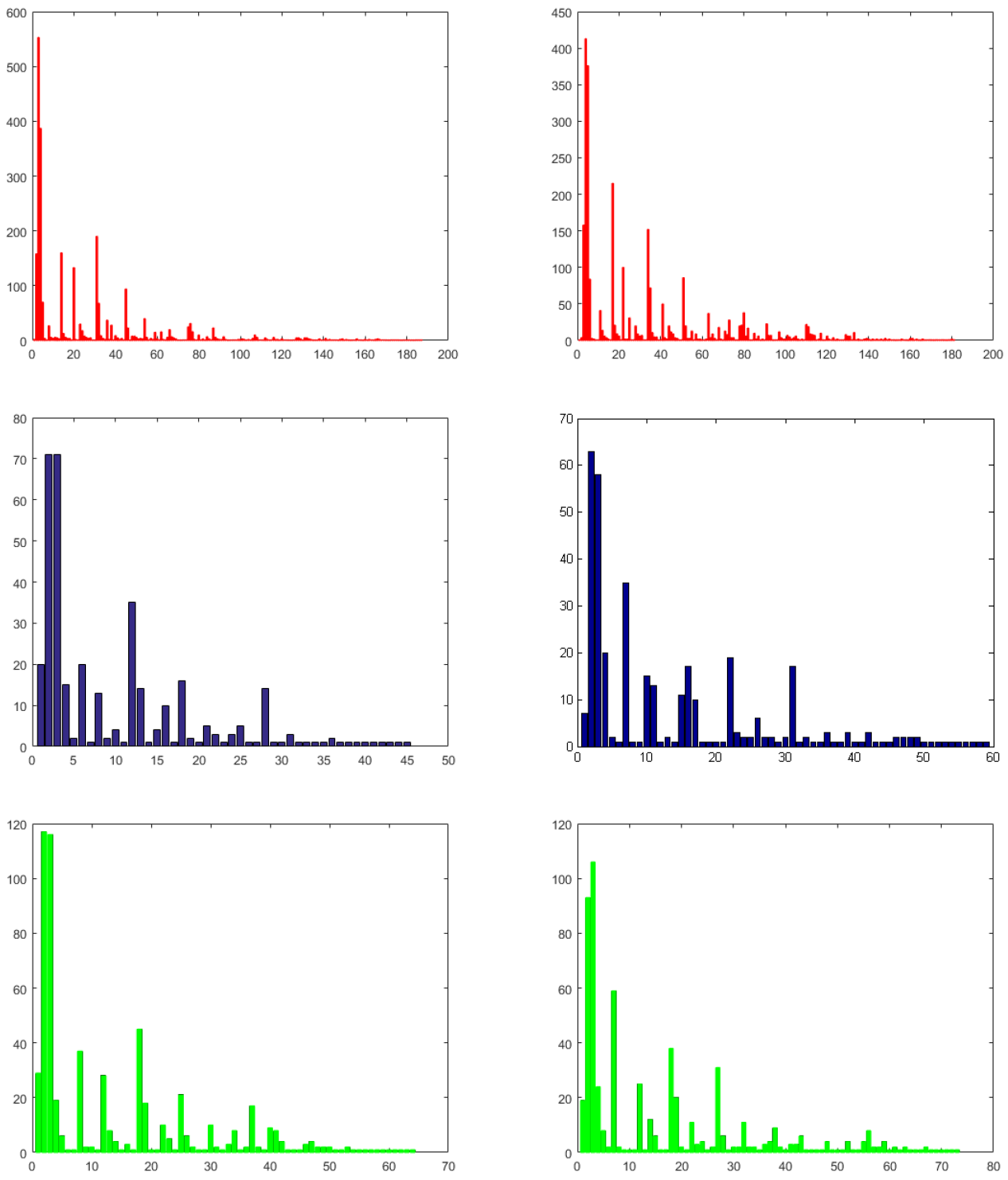
Bar plots of purine (on top) and pyrimidine (on bottom) distances of miRNAs of Human, Gorilla and Chimpanzee from left to right respectively.

It is found that most of the miRNAs of human, gorilla and chimpanzee have purine distance patterns [1 2 3] and [1 2 3 4]. There are exactly 553 and 387 many miRNAs of human belong to the cluster 3 ([1 2 3]) and 4 ([1 2 3 4]) respectively. In the case of miRNAs of gorilla, there are 71 miRNAs having purine distance pattern [1 2 3] and 71 miRNAs with distance pattern [1 2 3 4]. For miRNAs of chimpanzee, there are 117 and 116 distance patterns of purine [1 2 3] and [1 2 3 4]. There is no miRNAs of gorilla and chimpanzee having purine distance pattern [1]. In all the three cases, it is noted that there are several clusters having only one member which means that the mirRNA of those clusters has unique purine distance pattern. In the similar fashion, for all the miRNAs of human, gorilla and chimpanzee, the distance between pyrimidine bases to the next immediate pyrimidine bases are found, which is tabulated in the *Additional file-5 (sheet-1,2,3)*. The maximum length of the purine and pyrimidine distance patterns is found to be 5 for all the three set of miRNAs of human, gorilla and chimpanzee except five miRNAs only for human.

We also have determined the frequency distribution of purine and pyrimidine bases of the miRNAs of human, gorilla and chimpanzee as presented in detail in the *Additional file-6*. The clusters based on the frequency distribution of purine is made and tabulated in Table 4. The histogram of frequency distribution of purine bases for the three species are given in Fig. 8.

**Table 4.** Clusters based on Frequency distribution (Purine) of miRNAs of Human, Gorilla and Chimpanzee.

| Cluster | Human |  | Gorilla |  | Chimpanzee |  |
| --- | --- | --- | --- | --- | --- | --- |
|  | Frequency | Center | Frequency | Center | Frequency | Center |
| 1 | 27 | 0.0952 | 5 | 0.177 | 4 | 0.0987 |
| 2 | 114 | 0.1905 | 5 | 0.259 | 8 | 0.1786 |
| 3 | 253 | 0.2857 | 31 | 0.341 | 14 | 0.2584 |
| 4 | 381 | 0.381 | 81 | 0.423 | 59 | 0.3382 |
| 5 | 575 | 0.4762 | 110 | 0.505 | 126 | 0.4181 |
| 6 | 628 | 0.5714 | 70 | 0.586 | 138 | 0.4979 |
| 7 | 340 | 0.6667 | 28 | 0.668 | 129 | 0.5777 |
| 8 | 208 | 0.7619 | 20 | 0.75 | 67 | 0.6576 |
| 9 | 57 | 0.8571 | 6 | 0.832 | 32 | 0.7374 |
| 10 | 5 | 0.9524 | 1 | 0.914 | 10 | 0.8172 |

**Figure 8.**
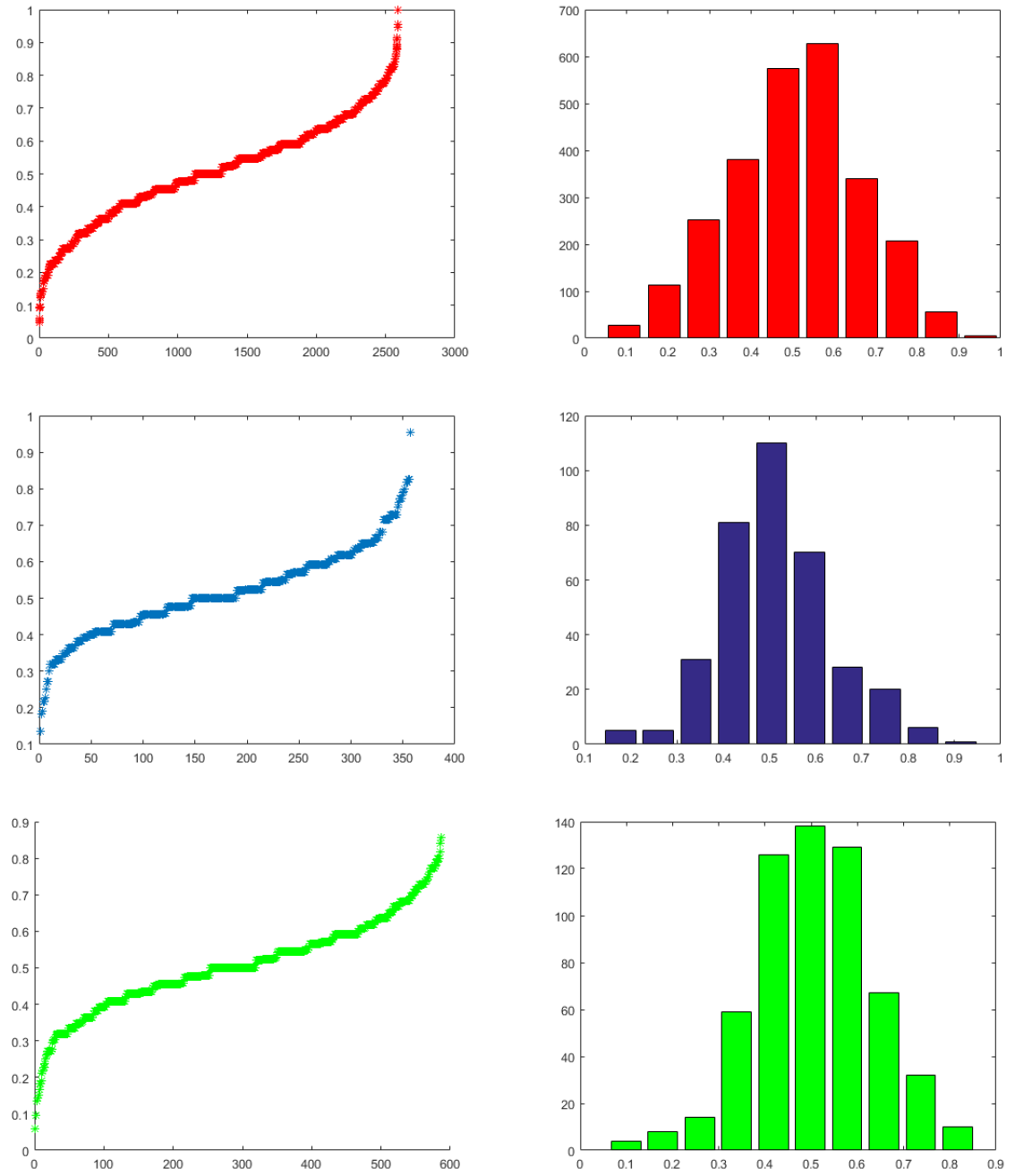
Histograms of frequency distribution of purine bases of the miRNAs of Human, Gorilla and Chimpanzee from top to bottom respectively.

**Figure 9.**
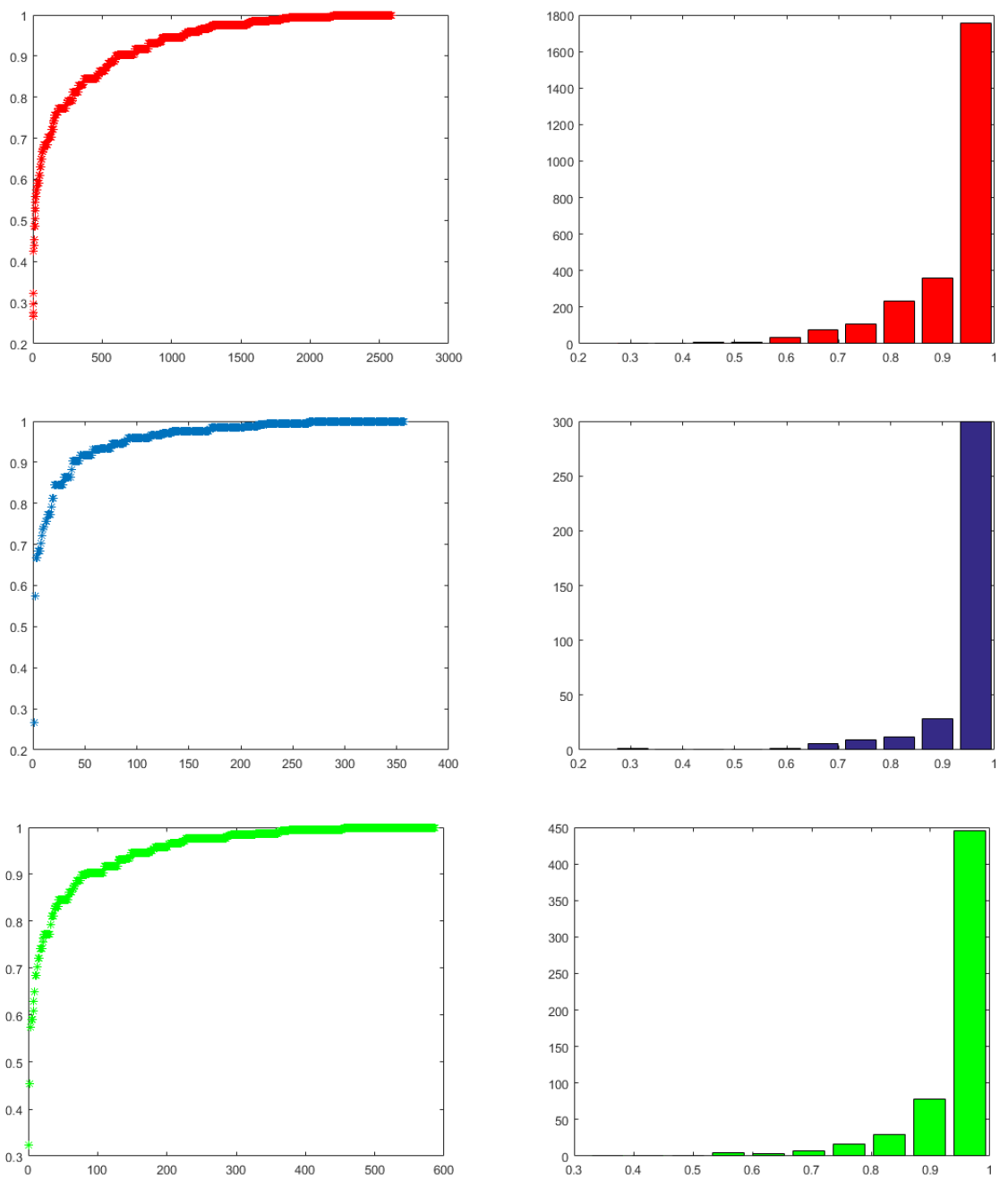
Histograms of Shannon entropy (purine and pyrimidine) Human, Gorilla and Chimpanzee from top to bottom respectively.

### Classification Based on SEs

For all the miRNAs of the three species human, gorilla and chimpanzee of the Hominidae family, the Shannon entropy is calculated of which detail can be seen in the *Additional file 7*. In the case of miRNAs of human, there are exactly 80 distinct SE values are obtained whereas in the case of miRNAs of gorilla and chimpanzee, there are 38 and 57 respectively distinct SEs are found. Based on the Shannon entropy, the miRNAs of the three species are classified into 10 clusters separately as shown in the Table 5. It is observed that most of the miRNAs of all the species human, gorilla and chimpanzee are having Shannon entropy centered at 0.96 which is the largest center of the clusters for all the three different clustering, which contain 1753, 300 and 446 miRNAs respectively. It is seen that there are 16 and 4 miRNAs of human and chimpanzee respectively centered at 0.5 (SE) but there is no member having same center 0.5 in the other set of miRNAs of gorilla. It is also seen that there is no miRNA of human and gorilla, whose Shannon entropy centered at 0.37 but there is only one miRNA of chimpanzee is having SE centered at 0.37. It interprets that none of the miRNAs of gorilla is having equal (approximately) purine-pyrimidine distribution over the sequences whereas there are 16 miRNAs of human which are approximately having random-like purine and pyrimidine distribution. Overall these observations draw an impression that almost none of the miRNAs of all three species is having random-like purine-pyrimidine distribution.

**Table 5.** Clusters based on Shannon entropy of miRNAs of Human, Gorilla and Chimpanzee.

| Cluster | Human |  | Gorilla |  | Chimpanzee |  |
| --- | --- | --- | --- | --- | --- | --- |
|  | Frequency | Center | Frequency | Center | Frequency | Center |
| 1 | 5 | 0.3034 | 1 | 0.3034 | 1 | 0.3566 |
| 2 | 0 | 0.3767 | 0 | 0.3767 | 1 | 0.4243 |
| 3 | 9 | 0.45 | 0 | 0.45 | 0 | 0.4921 |
| 4 | 7 | 0.5233 | 0 | 0.5233 | 4 | 0.5598 |
| 5 | 35 | 0.5967 | 1 | 0.5967 | 3 | 0.6275 |
| 6 | 76 | 0.67 | 6 | 0.67 | 7 | 0.6952 |
| 7 | 109 | 0.7433 | 9 | 0.7433 | 17 | 0.763 |
| 8 | 233 | 0.8167 | 12 | 0.8167 | 30 | 0.8307 |
| 9 | 360 | 0.89 | 28 | 0.89 | 78 | 0.8984 |
| 10 | 1753 | 0.9633 | 300 | 0.9633 | 446 | 0.9661 |

## Discussion

The purine and pyrimidine analysis with the binomial distribution shows the purines and pyrimidines are not independently distributed over the miRNAs and there is a tendency of same properties (purine or pyrimidine) to repeat in a miRNA. We found the various classes using different methods where most of the cases the classes are normally distributed although the distribution of the purines and pyrimidines is not random like distribution.

There are 5 miRNAs of human in the cluster 10 based on fractal dimension as shown in the Table-1 having maximum FD. We have seen closely those sequences and found that three of them (h2248, h1954 and h2552) are pyrimidine rich sequences (94%, 90% and 90% respectively) and other two (h1835 and h1291) are purine rich sequence (95% and 100 %) as shown in Table-6. In the case of miRNAs of gorilla and chimpanzee, the cluster 10 contains 2 and 3 miRNAs respectively. Here we convict that whenever the amount of purine or pyrimidine is highly rich in the miRNA sequence then corresponding FD will be maximum.

**Table 6.**
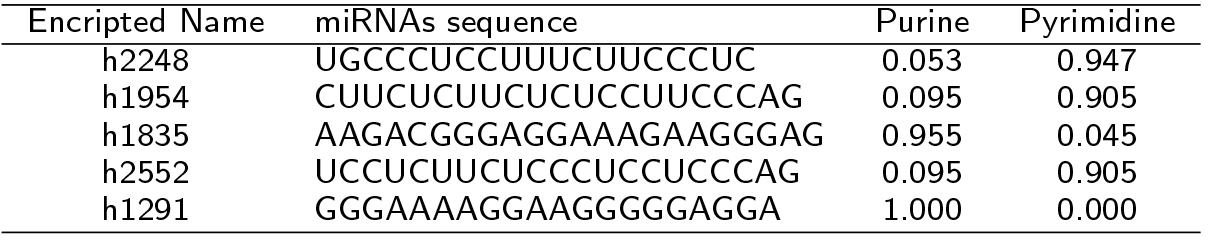
MiRNAs (from the cluster-10 based on fractal dimension) and its frequency distribution of miRNAs of Human.

There are several clusters having miRNAs for human, gorilla and chimpanzee with the same HEs. The frequency distribution of purine and pyrimidine are balanced for all such miRNAs having same HEs. For an example, we took 17 miRNAs from the cluster 7 of miRNAs of human based on Hurst exponent (h463, h526, ..., h1824 and h2202) which all are having same HE (0.714484) as shown in the Table -7. It is found that the frequency distribution of purine and pyrimidine 60% and 40% (or 40% and 60%) respectively. It is also seen that these 17 miRNAs of human belong to same cluster 3 based on fractal dimension. It is observed that there are 6% miRNAs of human and chimpanzee and 14% miRNAs of gorilla having HE 0.5 which indicates a completely uncorrelated purine and pyrimidine spatial ordering over the miRNAs.

**Table 7.**
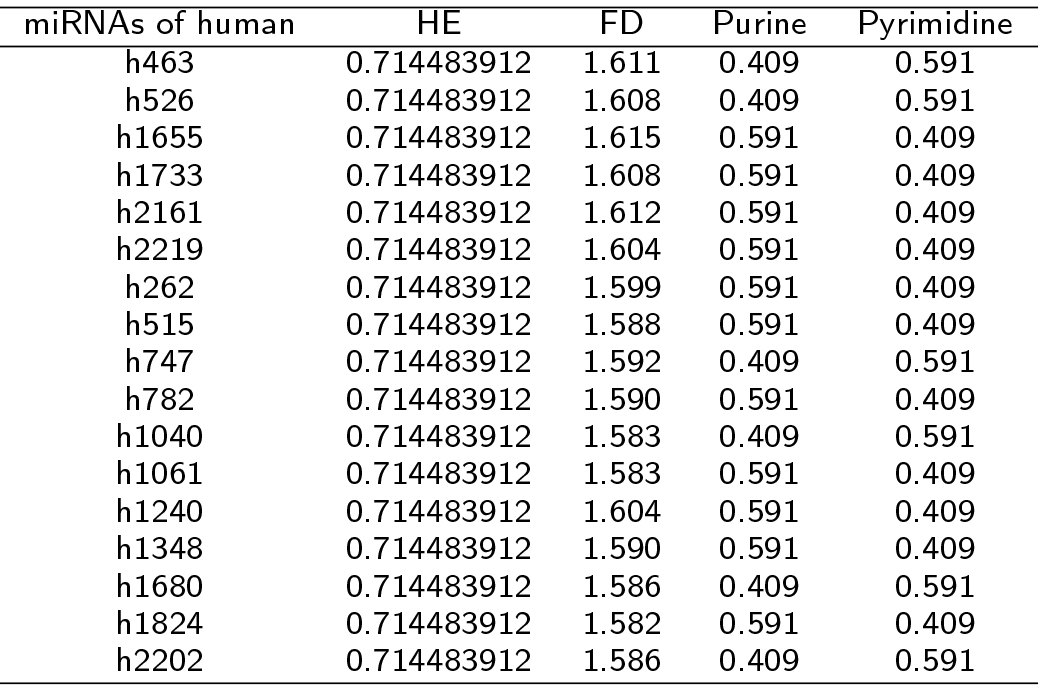
MiRNAs (from the cluster-7 based on Hurst exponent) and its FD with the frequency distribution of miRNAs of Human.

| miRNAs of human | HE | FD | Purine | Pyrimidine |
| --- | --- | --- | --- | --- |
| h463 | 0.714483912 | 1.611 | 0.409 | 0.591 |
| h526 | 0.714483912 | 1.608 | 0.409 | 0.591 |
| h1655 | 0.714483912 | 1.615 | 0.591 | 0.409 |
| h1733 | 0.714483912 | 1.608 | 0.591 | 0.409 |
| h2161 | 0.714483912 | 1.612 | 0.591 | 0.409 |
| h2219 | 0.714483912 | 1.604 | 0.591 | 0.409 |
| h262 | 0.714483912 | 1.599 | 0.591 | 0.409 |
| h515 | 0.714483912 | 1.588 | 0.591 | 0.409 |
| h747 | 0.714483912 | 1.592 | 0.409 | 0.591 |
| h782 | 0.714483912 | 1.590 | 0.591 | 0.409 |
| h1040 | 0.714483912 | 1.583 | 0.409 | 0.591 |
| h1061 | 0.714483912 | 1.583 | 0.591 | 0.409 |
| h1240 | 0.714483912 | 1.604 | 0.591 | 0.409 |
| h1348 | 0.714483912 | 1.590 | 0.591 | 0.409 |
| h1680 | 0.714483912 | 1.586 | 0.409 | 0.591 |
| h1824 | 0.714483912 | 1.582 | 0.591 | 0.409 |
| h2202 | 0.714483912 | 1.586 | 0.409 | 0.591 |

Now we shall see the miRNAs of human having identical distance pattern of purine and pyrimidine. It is observed that there are very few numbers of miRNAs of human which are having identical distance pattern of purine and pyrimidine. For example, h61 miRNA is having identical distance pattern of purine and pyrimidine [1 2]. Also There are miRNAs h36, h51, h62, h83, h122 and h2584 having identical distance pattern of purine and pyrimidine [1 2 3]. There are miRNAs of gorilla g6, g13, g19, g20 and g21 having identical distance pattern of purine and pyrimidine [1 2 3 4]. In the set of miRNAs of chimpanzee, there are p55, p84, p106, p119 and p138 having identical purine and pyrimidine distance pattern. This identical distance pattern of purine and pyrimidine make a guarantee that there are purine and pyrimidine block of same length.

There are 142 distinct frequency distribution of purine and pyrimidine bases across miRNAs of human are found. Out of 2588 miRNAs of human, 194 miRNAs of human having equal frequency distribution of purine and pyrimidine and 1121 miRNAs of human having lesser frequency of purines than that of pyrimidine of 1273 miRNAs are there. This infers frequency of pyrimidine is richer than that of purine over the set of miRNAs of human. On the other hand, there are 43 miRNAs out of 357 miRNAs of gorilla having equal frequency distribution of purine and pyrimidine. There are 58 distinct frequency distribution of purine and pyrimidine bases across miRNAs of gorilla. Out of 357 miRNAs of gorilla, 146 miRNAs of gorilla having lesser frequency of purines than that of pyrimidine of 168 miRNAs are there. There are 92 distinct frequency distribution of purine and pyrimidine bases across miRNAs of chimpanzee are found. Out of 587 miRNAs of human, 67 miRNAs of chimpanzee having equal frequency distribution of purine and pyrimidine and 268 miRNAs of human having lesser frequency of purines than that of pyrimidine of 252 miRNAs are there. This infers frequency of pyrimidine is richer than that of purine over the set of miRNAs of chimpanzee. In this regard we infer that the frequency distribution of pyrimidine over the miRNAs of these three observed species human, gorilla and chimpanzee is richer than that of the purine bases. Out of all the 2588 miRNAs, the miRNA h1291 is only miRNA containing all purine bases.

It is reported that MiR-200 is a family of tumour suppressor miRNAs consisting of five members (h609, h888, h1449, h1677, h1520, h2186, h515, h328 and h670), which are significantly involved in inhibition of epithelial-to-mesenchymal transition (EMT), repression of cancer stem cells (CSCs) self-renewal and differentiation, modulation of cell division and apoptosis, and reversal of chemoresistance [18] as shown in *Table-8*. We have chosen all these nine miRNAs of human including other miRNAs of human which are 0, 1 and 2 Hamming distance apart from those nine miRNAs.

**Table 8.** MiRNAs Star miRNAs in human cancer and their quantifications.

| HD | miRNAs | FD | HE | Freq. of Purine | Freq. of Pyrimidine |
| --- | --- | --- | --- | --- | --- |
|  | **h609** | **1.579** | **0.578** | **0.5** | **0.5** |
| HD=1 | h888 | 1.580 | 0.649 | 0.545 | 0.455 |
| HD=2 | h515 | 1.588 | 0.714 | 0.591 | 0.409 |
| HD=2 | h875 | 1.587 | 0.610 | 0.478 | 0.522 |
| HD=2 | h1520 | 1.604 | 0.627 | 0.609 | 0.391 |
| HD=2 | h1670 | 1.579 | 0.5 | 0.5 | 0.5 |
|  | **h888** | **1.580** | **0.649** | **0.545** | **0.455** |
| HD=1 | h515 | 1.588 | 0.714 | 0.5 | 0.5 |
| HD=1 | h609 | 1.579 | 0.578 | 0.5 | 0.5 |
| HD=2 | h1236 | 1.569 | 0.654 | 0.647 | 0.353 |
| HD=2 | h1520 | 1.604 | 0.627 | 0.609 | 0.391 |
|  | **h1449** | **1.579** | **0.671** | **0.5** | **0.5** |
| HD=0 | h1677 | 1.579 | 0.671 | 0.5 | 0.5 |
| HD=2 | h749 | 1.509 | 0.655 | 0.438 | 0.563 |
| HD=2 | h2186 | 1.596 | 0.638 | 0.409 | 0.591 |
|  | **h1677** | **1.579** | **0.671** | **0.545** | **0.455** |
| HD=0 | h1449 | 1.579 | 0.671 | 0.5 | 0.5 |
| HD=2 | h749 | 1.509 | 0.655 | 0.438 | 0.563 |
| HD=2 | h2186 | 1.601 | 0.638 | 0.409 | 0.591 |
|  | **h1520** | **1.604** | **0.627** | **0.609** | **0.391** |
| Hd=1 | h888 | 1.580 | 0.649 | 0.545 | 0.455 |
| HD=2 | h515 | 1.588 | 0.714 | 0.591 | 0.409 |
| HD=2 | h609 | 1.579 | 0.578 | 0.5 | 0.5 |
| HD=2 | h2413 | 1.590 | 0.528 | 0.706 | 0.294 |
|  | **h2186** | **1.596** | **0.638** | **0.409** | **0.591** |
| Hd=1 | h749 | 1.509 | 0.655 | 0.438 | 0.563 |
| HD=2 | h1449 | 1.579 | 0.671 | 0.5 | 0.5 |
| HD=2 | h1677 | 1.579 | 0.671 | 0.545 | 0.455 |
|  | **h515** | **1.588** | **0.714** | **0.591** | **0.409** |
| Hd=1 | h888 | 1.580 | 0.649 | 0.545 | 0.455 |
| HD=1 | h1236 | 1.569 | 0.654 | 0.647 | 0.353 |
| HD=2 | h609 | 1.579 | 0.578 | 0.5 | 0.5 |
| HD=2 | h712 | 1.543 | 0.745 | 0.556 | 0.444 |
| HD=2 | h782 | 1.590 | 0.714 | 0.591 | 0.409 |
| HD=2 | h1520 | 1.604 | 0.627 | 0.609 | 0.391 |
| HD=2 | h1670 | 1.579 | 0.671 | 0.5 | 0.5 |
|  | **h328** | **1.585** | **0.709** | **0.455** | **0.545** |
| HD=0 | × |  |  |  |  |
| HD=1 | × |  |  |  |  |
| HD=2 | × |  |  |  |  |
|  | **h1670** | **1.579** | **0.671** | **0.5** | **0.5** |
| Hd=2 | h515 | 1.588 | 0.714 | 0.591 | 0.409 |
| HD=2 | h609 | 1.579 | 0.578 | 0.5 | 0.5 |
| HD=2 | h2485 | 1.533 | 0.873 | 0.471 | 0.529 |

It is found that h888 is 1 Hamming distance apart from h609 and h1520 although h609 and h1520 are 2 Hamming distance apart. The miRNA h888 is having approximately same HD, HE and frequencies of purine and pyrimidine bases with the miRNAs h609 and h1520. Hence we convict that h888 might also work as h609 and h1520 do. It is also observed that h1670 is 2 HD apart from the miRNAs h609 and h515 and the miRNA h1670 is showing very close as per quantitative measures and hence this miRNA h1670 would function as h609 and h515 do. There are two miRNAs of human h1449 and h1677 are 0 Hamming distance apart with same quantitative measures and hence we firmly propose that these two miRNAs would functions same. It is worth noting that both the miRNAs h1449 and h1677 have identical purine-pyrimidine organization. Following the similar argument, other association with the rest of miRNAs can also be made. It is seen that there does not exist any human miRNA which is 0, 1 or 2 HD apart from the miRNA h328.

## Concluding Remarks

One of the integral divisions of nucleotides based on their chemical properties is purine-pyrimidine. We attempted to understand the distribution of purine and pyrimidine bases over all the miRNAs in three species human, gorilla and chimpanzee. Quantitatively, we deciphered the self-organization of the purine and pyrimidine bases for all the miRNAs through the fractal dimension of the indicator matrix. Also we took out the auto correlation of purine-pyrimidine bases through the parameter Hurst exponent. To get the nearness of the miRNAs based on their purine-pyrimidine distribution, HD is deployed. The purine-pyrimidine distance patterns including the frequency distribution have been found for all the miRNAs. For all these parameters, we did cluster the miRNAs into several clusters. Based on the quantitative investigation, some crucial observations are adumbrated in the discussion. It is worth noting that through our investigation, the miRNA h1291, made of only purine bases is identified.

Through these investigations what it fails to attain is the association among the miRNAs and target mRNAs as shown in *Table 9*, *Table 10* and *Table 11*. The complex relationship among miRNAs and target mRNAs, previously pointed out in literatures [12] has been reconfirmed through our quantitative analysis over a set of miRNAs and target mRNAs. Imperfect base-pairing between the miRNA and the 3’-UTR of its target mRNA leads to blockage of translation, or at least accumulation of the mRNA’s protein product, whereas perfect or near-perfect base-pairing between the miRNA and the middle of its target mRNA causes cleavage of the mRNA, thereby inactivating the same. Such diverse patterns of miRNA action may be responsible for making the correlation among miRNA and target mRNA complex, that are yet to be resolved decisively as pointed out by several researchers earlier. In this context, we plan to bring out the pattern of organization of nucleotides following the other two modes of classifications (amino-keto and strong H-bond and weak H-bond) based on the chemical properties of nucleotides in order to integrate the whole three kinds of grouping to find out the correlationship among the miRNAs-mRNAs and corresponding targeted regions of mRNAs.

**Table 9.**
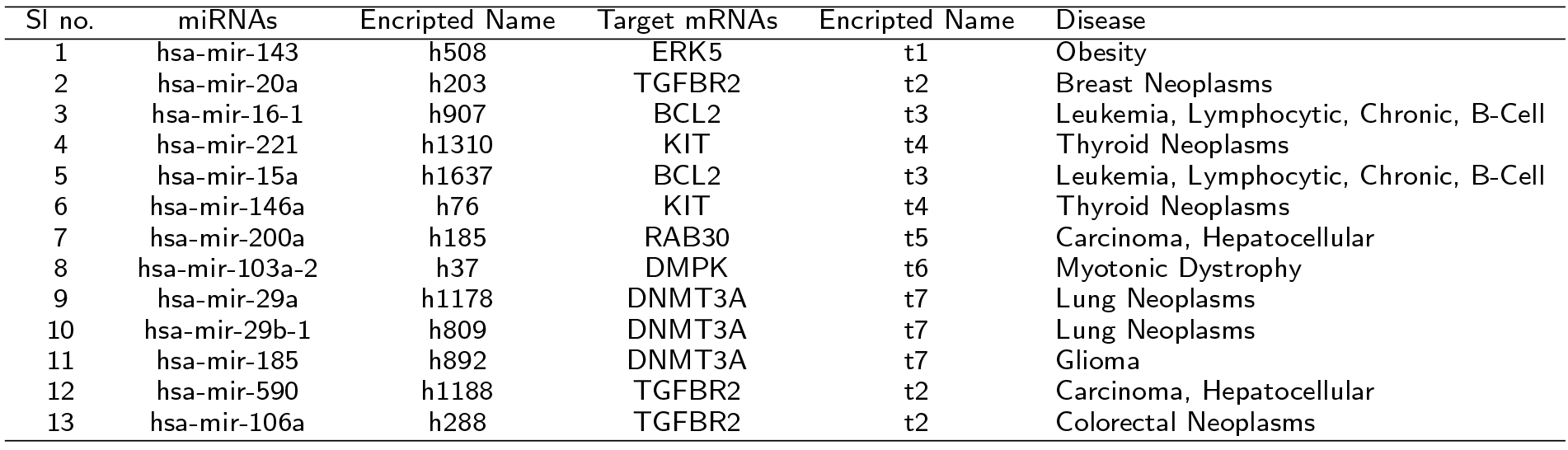
Selected miRNAs of Human and the corresponding target mRNAs.

| Sl no. | miRNAs | Encripted Name | Target mRNAs | Encripted Name | Disease |
| --- | --- | --- | --- | --- | --- |
| 1 | hsa-mir-143 | h508 | ERK5 | t1 | Obesity |
| 2 | hsa-mir-20a | h203 | TGFBR2 | t2 | Breast Neoplasms |
| 3 | hsa-mir-16-1 | h907 | BCL2 | t3 | Leukemia, Lymphocytic, Chronic, B-Cell |
| 4 | hsa-mir-221 | h1310 | KIT | t4 | Thyroid Neoplasms |
| 5 | hsa-mir-15a | h1637 | BCL2 | t3 | Leukemia, Lymphocytic, Chronic, B-Cell |
| 6 | hsa-mir-146a | h76 | KIT | t4 | Thyroid Neoplasms |
| 7 | hsa-mir-200a | h185 | RAB30 | t5 | Carcinoma, Hepatocellular |
| 8 | hsa-mir-103a-2 | h37 | DMPK | t6 | Myotonic Dystrophy |
| 9 | hsa-mir-29a | h1178 | DNMT3A | t7 | Lung Neoplasms |
| 10 | hsa-mir-29b-1 | h809 | DNMT3A | t7 | Lung Neoplasms |
| 11 | hsa-mir-185 | h892 | DNMT3A | t7 | Glioma |
| 12 | hsa-mir-590 | h1188 | TGFBR2 | t2 | Carcinoma, Hepatocellular |
| 13 | hsa-mir-106a | h288 | TGFBR2 | t2 | Colorectal Neoplasms |

**Table 10.**

| Target mRNAs | FD | HE | Purine distance pattern | Pyrimidine distance pattern |
| --- | --- | --- | --- | --- |
| t1 | 1.94 | 0.6634044 | [1 2 3 4 5 6 7 8 9 10 11] | [1 2 3 4 5 6 7 8 9 10 12 13] |
| t2 | 1.89362 | 0.5792666 | [1 2 3 4 5 6 7 8 9 10] | [1 2 3 4 5 6 7 8 10 11 17] |
| t3 | 1.8819 | 0.6976582 | [1 2 3 4 5 6 7 8 9 11 13 17] | [1 2 3 4 5 6 7 8 9 10 11] |
| t4 | 1.92328 | 0.6255126 | [1 2 3 4 5 6 7 8 9 10 12 13 15] | [1 2 3 4 5 6 7 8 9 10 11 12 14 16 18] |
| t5 | 1.8773 | 0.5928585 | [1 2 3 4 5 6 7 8 10 13] | [1 2 3 4 5 6 7 8 9 14 15] |
| t6 | 1.90762 | 0.7073991 | [1 2 3 4 5 6 7 8 9 10 12 13 14] | [1 2 3 4 5 6 7 8 9 10 12 14 19] |
| t7 | 1.91941 | 0.6735526 | [1 2 3 4 5 6 7 8 9 10 11 12] | [1 2 3 4 5 6 7 8 9 10 11 12 13 16] |

**Table 11.**
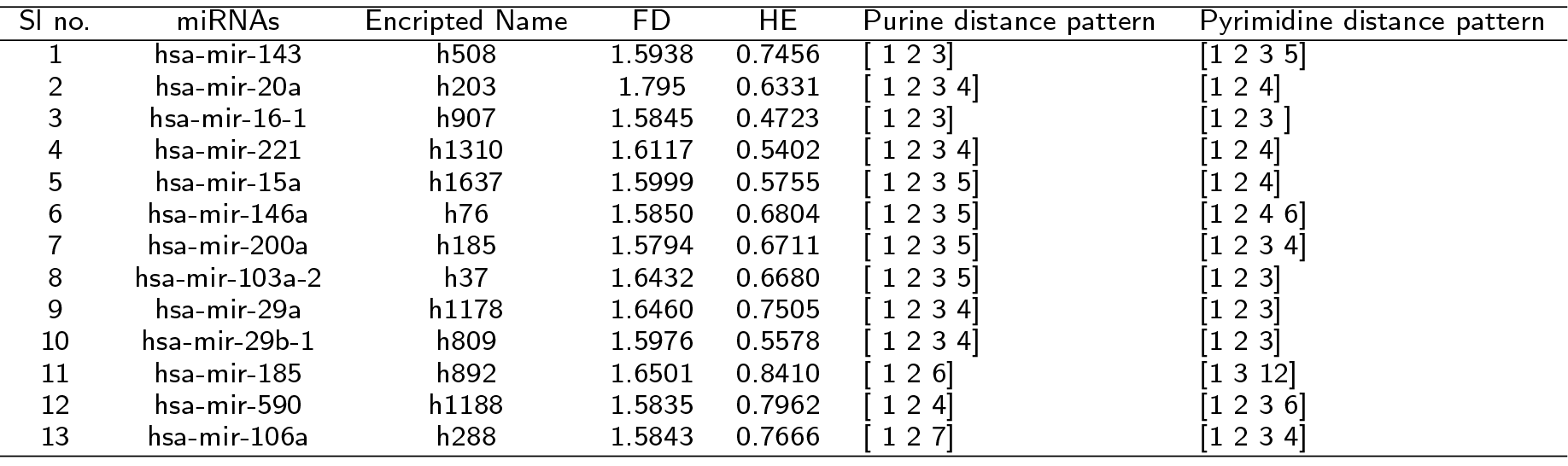
MiRNAs of Human and Corresponding Quantifications

| Sl no. | miRNAs | Encripted Name | FD | HE | Purine distance pattern | Pyrimidine distance pattern |
| --- | --- | --- | --- | --- | --- | --- |
| 1 | hsa-mir-143 | h508 | 1.5938 | 0.7456 | [1 2 3] | [1 2 3 5] |
| 2 | hsa-mir-20a | h203 | 1.795 | 0.6331 | [1 2 3 4] | [1 2 4] |
| 3 | hsa-mir-16-1 | h907 | 1.5845 | 0.4723 | [1 2 3] | [1 2 3] |
| 4 | hsa-mir-221 | h1310 | 1.6117 | 0.5402 | [1 2 3 4] | [1 2 4] |
| 5 | hsa-mir-15a | h1637 | 1.5999 | 0.5755 | [1 2 3 5] | [1 2 4] |
| 6 | hsa-mir-146a | h76 | 1.5850 | 0.6804 | [1 2 3 5] | [1 2 4 6] |
| 7 | hsa-mir-200a | h185 | 1.5794 | 0.6711 | [1 2 3 5] | [1 2 3 4] |
| 8 | hsa-mir-103a-2 | h37 | 1.6432 | 0.6680 | [1 2 3 5] | [1 2 3] |
| 9 | hsa-mir-29a | h1178 | 1.6460 | 0.7505 | [1 2 3 4] | [1 2 3] |
| 10 | hsa-mir-29b-1 | h809 | 1.5976 | 0.5578 | [1 2 3 4] | [1 2 3] |
| 11 | hsa-mir-185 | h892 | 1.6501 | 0.8410 | [1 2 6] | [1 3 12] |
| 12 | hsa-mir-590 | h1188 | 1.5835 | 0.7962 | [1 2 4] | [1 2 3 6] |
| 13 | hsa-mir-106a | h288 | 1.5843 | 0.7666 | [1 2 7] | [1 2 3 4] |

## Author’s contributions

PPC, SSH and AC conceptualize the problem and JKD, SSH and PB performs computational analysis of the datasets. SSH and JKD wrote the manuscript and finally edited and corrected by AC, PB and PPC.

## Acknowledgments

The authors acknowledge Prof. R. L. Brahmachary for his valuable suggestions.

## Additional Files

Additional file 1 – Name Encryption of miRNA of Human, Gorilla and Chimpanzee.

Additional file 2 – Fractal Dimension (FD) of all binary sequences of Human, Gorilla and Chimpanzee (sheet-1) and the detail members (miRNAs) of the clusters for Human, Gorilla and Chimpanzee based on FD (sheet-2).

Additional file 3 – Hurst Exponent (HE) of all binary sequences of Human, Gorilla and Chimpanzee (sheet-1) and The detail members (miRNAs) of the clusters for Human, Gorilla and Chimpanzee based on HE (sheet-2).

Additional file 4 – The Hamming Distance (HD) matrices of (miRNAs) of Human (sheet 1), Gorilla (sheet 2) and Chimpanzee (sheet 3) between the pair of two sequences which can be identified by the corresponding row and column numbers.

Additional file 5 – The detail clustering of miRNAs of Human (sheet 1), Gorilla (sheet 2) and Chimpanzee (sheet 3) with cluster numbers based on the immediate purine and pyrimidine distances as patterns.

Additional file 6 – The detail frequency distribution of purine and pyrimidine of miRNAs of Human, Gorilla and Chimpanzee.

Additional file 7 – Shannon Entropy (SE) of all binary sequences of Human, Gorilla and Chimpanzee (sheet-1 and the detail members (miRNAs) of the clusters for Human, Gorilla and Chimpanzee based on SE (sheet-2).

